# Analysis of the Genomic Basis of Functional Diversity in Dinoflagellates using a Transcriptome-Based Sequence Similarity Network

**DOI:** 10.1101/211243

**Authors:** Arnaud Meng, Erwan Corre, Ian Probert, Andres Gutierrez-Rodriguez, Raffaele Siano, Anita Annamale, Adriana Alberti, Corinne Da Silva, Patrick Wincker, Stéphane Le Crom, Fabrice Not, Lucie Bittner

## Abstract

Dinoflagellates are one of the most abundant and functionally diverse groups of eukaryotes. Despite an overall scarcity of genomic information for dinoflagellates, constantly emerging high-throughput sequencing resources can be used to characterize and compare these organisms. We assembled *de novo* and processed 46 dinoflagellate transcriptomes and used a sequence similarity network (SSN) to compare the underlying genomic basis of functional features within the group. This approach constitutes the most comprehensive picture to date of the genomic potential of dinoflagellates. A core proteome composed of 252 connected components (CCs) of putative conserved protein domains (pCDs) was identified. Of these, 206 were novel and 16 lacked any functional annotation in public databases. Integration of functional information in our network analyses allowed investigation of pCDs specifically associated to functional traits. With respect to toxicity, sequences homologous to those of proteins involved in toxin biosynthesis pathways (e.g. *sxtA1-4* and *sxtG*) were not specific to known toxin-producing species. Although not fully specific to symbiosis, the most represented functions associated with proteins involved in the symbiotic trait were related to membrane processes and ion transport. Overall, our SSN approach led to identification of 45,207 and 90,794 specific and constitutive pCDs of respectively the toxic and symbiotic species represented in our analyses. Of these, 56% and 57% respectively (*i.e.* 25,393 and 52,193 pCDs) completely lacked annotation in public databases. This stresses the extent of our lack of knowledge, while emphasizing the potential of SSNs to identify candidate pCDs for further functional genomic characterization.

## INTRODUCTION

Dinoflagellates are unicellular eukaryotes belonging to the Alveolata lineage (Bachvaroff *et al.* 2014). This group encompasses a broad diversity of taxa that have a long and complex evolutionary history, play key ecological roles in aquatic ecosystems, and have significant economic impacts (Murray *et al.* 2016; Janouškovec *et al.* 2016). The ecological success of dinoflagellates in the marine planktonic environment is assumed to be due to their ability to exhibit various survival strategies associated with an extraordinary physiological diversity (Murray *et al.* 2016). Nearly half of dinoflagellates have chloroplasts, but most of these are likely mixotrophic, combining photosynthetic and heterotrophic modes of nutrition (Jeong *et al.* 2010; Stoecker *et al.* 2017). Many dinoflagellates produce toxins and form long-lasting harmful algal blooms with deleterious effects on fisheries or aquaculture (Flewelling *et al.* 2005). Some species of the genus *Alexandrium* can produce toxins that effect higher trophic levels in marine ecosystems (*i.e.* copepods, fish) and are harmful to humans (Orr *et al.* 2013; Kohli *et al.* 2016; Murray *et al.* 2016). Members of the genus *Symbiodinium* are known to establish mutualistic symbioses with a wide diversity of benthic hosts, sustaining reef ecosystems worldwide (Goodson *et al.* 2001; Lin *et al.* 2015). Interactions between dinoflagellates and other marine organisms are extremely diverse, including (photo)symbioses (Decelle *et al.* 2015), predation (Jeong *et al.* 2010), kleptoplasty (Gast *et al.* 2007), and parasitism (Siano *et al.* 2011). Dinoflagellates have been highlighted as important members of coastal and open-ocean protistan communities based on environmental molecular barcoding surveys (Massana *et al.* 2015; Le Bescot *et al.* 2016) and the parasitic syndiniales in particular have been identified as key players that drive *in situ* planktonic interactions in the ocean (Lima-Mendez *et al.* 2015).

Along with metabarcoding surveys based on taxonomic marker genes, environmental investigations of protistan ecology and evolution involve genomic and transcriptomic data. Interpretation of such large datasets is limited by the current lack of reference data from unicellular eukaryotic planktonic organisms, resulting in a high proportion of unknown sequences (Caron *et al.* 2016; Sibbald & Archibald 2017). This is particularly significant for dinoflagellates as this taxon remains poorly explored at the genome level, with only three full genome sequences published so far (Shoguchi *et al.* 2013; Lin *et al.* 2015; Aranda *et al.* 2016). Their genomes are notoriously big (0.5 to 40x larger than the human haploid genome) and have a complex organization (Jaeckisch *et al.* 2011; Shoguchi *et al.* 2013; Murray *et al.* 2016). Consequently, most recent studies investigating functional diversity of dinoflagellates rely on transcriptomic data to probe these non-model organisms.

The Moore Foundation Marine Microbial Eukaryotic Transcriptome Sequencing Project (MMETSP, http://marinemicroeukaryotes.org/, (Keeling *et al.* 2014)) provided the opportunity to produce a large quantity of reference transcriptomic data (Sibbald & Archibald 2017). Among the 650 transcriptomes released, 56 were from 24 dinoflagellate genera encompassing 46 distinct strains (Keeling *et al.* 2014). This dataset constitutes a unique opportunity to investigate the genomic basis of the major evolutionary and ecological traits of dinoflagellates (Janouškovec *et al.* 2016). Performing a global analysis of such a large dataset (~3 million sequences) is challenging and requires innovative approaches. Most studies published so far have targeted specific biological processes and pathways, focusing on a small subset of the available data (Meyer *et al.* 2015; Dupont *et al.* 2015; Kohli *et al.* 2016). In one recent study a 101-protein dataset was used to produce a multiprotein phylogeny of dinoflagellates (Janouškovec *et al.* 2016). As a large fraction of the sequences produced in the MMETSP project do not have any distant homologues in current reference databases, almost half (46%) of the data remains unannotated.

With the advent of high-throughput sequencing technologies and its inherent massive production of data, sequence similarity network (SSN) approaches (Atkinson *et al.* 2009; Cheng *et al.* 2014; Méheust *et al.* 2016) offer an alternative to classical methods, enabling inclusion of unknown sequences in the global analysis (Forster *et al.* 2015; Lopez *et al.* 2015). In a functional genomic context, SSNs facilitate large-scale comparison of sequences, including functionally unannotated sequences, and hypothesis design based on both model and non-model organisms. For instance, SSN has been used to define enolase protein superfamilies and assign function to nearly 50% of sequences composing the superfamilies that had unknown functions (Gerlt *et al.* 2012). Here we used a SSN approach involving 57 *de novo* assembled transcriptomes from the MMETSP project as well as new transcriptomes of four recently described dinoflagellates to unveil the core-, accessory-, and pan-proteome of dinoflagellates and to define gene sets characteristic of selected functional traits.

## RESULTS

### Dataset metrics overview

For the 57 assembled transcriptomes (53 from the MMETSP dataset and 4 from this study) the average number of transcript sequences was 93,685 with a mean N50 of 878 bp and remapping rates exceeded 60% on average (Tab. 1). Protein coding domains were predicted from the transcript sequences for each transcriptome to build what we consider here as proteomes. We found a mean of 49,281 protein-coding domains per proteome (Tab. 1). Globally, more than half of the protein-coding domains matched with functional annotations in InterPro (58%: 750,480 of 1,283,775) of which 552,846 had an identified Gene Ontology (GO) annotation. All individually assembled transcriptomes, derived proteomes and their corresponding functional annotations are available at http://application.sb-roscoff.fr/project/radiolaria/.

**Tab. 1:**
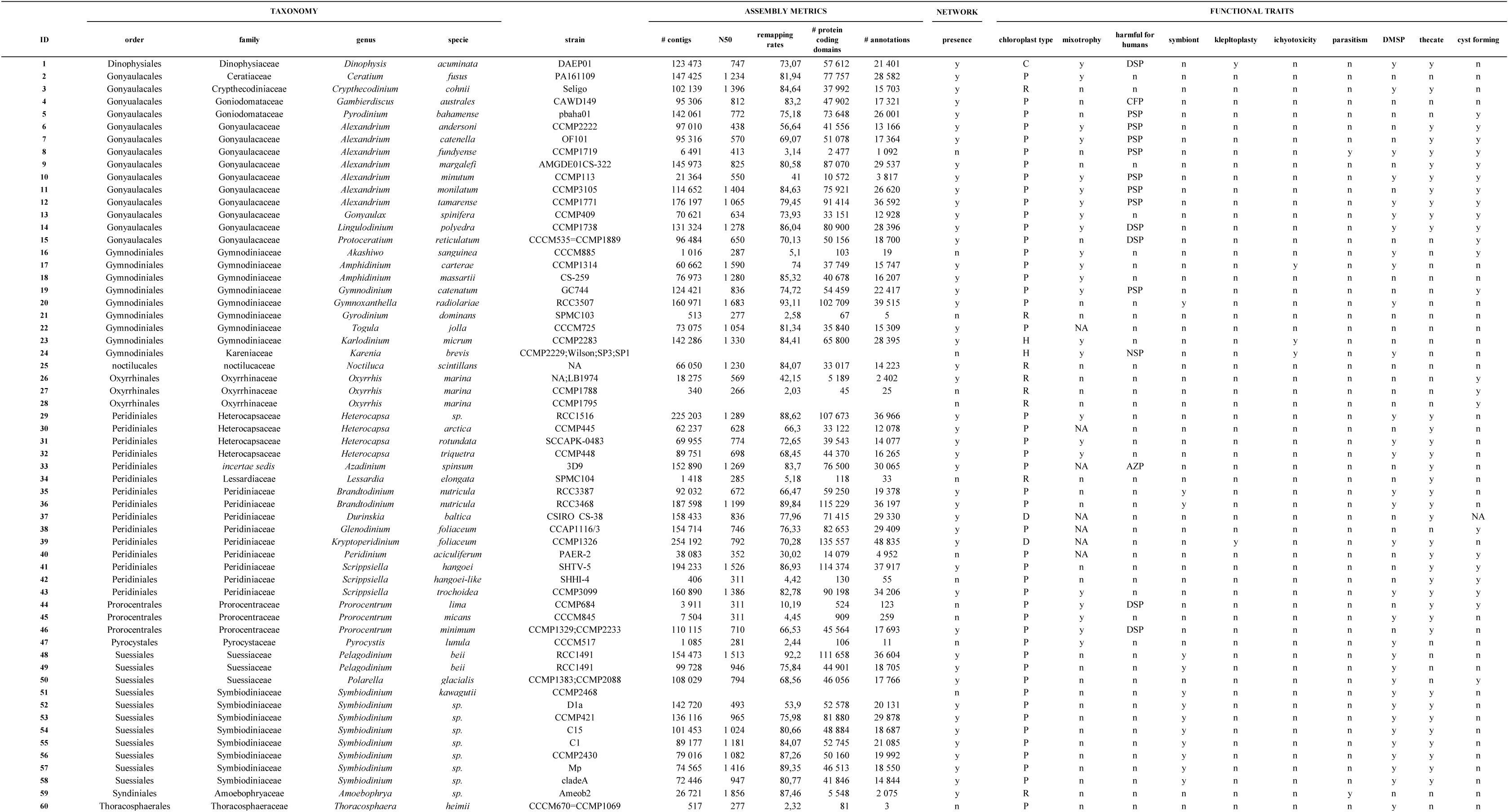
Summary table of the 60 transcriptomes (and the corresponding strains) analyzed in this study, ranked based on their taxonomy. Assembly metrics are reported for each transcriptome encompassing: the number of assembled contigs, N50, the remapping rate of initial reads, the number of predicted protein domains found in transcript sequences and the number of functional annotations identified through Interproscan 5. The network presence column indicates the “high quality” transcriptomes for which derived proteomes were included in the final network. Based on a literature survey, information about functional traits for each species included in the dataset is provided: chloroplast type (P: peridinin, H: haptophyte-like, C: cryptomonad-like, D: diatom-like and R: remnant or absent plastid), mixotrophy, ability to produce toxins harmful for humans (DSP:Diarrhetic shellfish poisoning, CFP:Ciguatera Fish Poisoning, PSP:Paralytic shellfish poisoning, AZP:Azaspiracid Shellfish Poisoning, NSP:Neurologic Shellfish Poisoning), ability to be symbionts, kleptoplasty, ichyotoxicity, parasitism, ability to produce DSMP, presence of a theca, ability to form cysts during life-cycle. <NA> corresponds to a lack of information.

A first version of the SSN was created based on the 57 proteomes. The filtration steps performed according to determined optimal settings (*i.e.* alignments with a minimum 60% sequence identity) (Fig. S1, see methods) resulted in the removal of 11 proteomes. The final network was composed of 1,275,911 vertices (protein-coding domains) linked by 6,142,013 edges (corresponding to a pairwise sequence identity value ≥ 60%). The network consisted of 350,267 connected components (CCs) with 11,568 of these having a size from 10 to 100 vertices (Tab. S1). It encompassed 46 proteomes having a mean of 60,661 protein-coding domains with an average length of 307 bp. According to InterPro functional annotations, 50.5% (*i.e.* 176,958) of the CCs were composed of unannotated sequences only.

### Identification of core / accessory / pan connected components

Analyses of CC composition revealed that 252 CCs included protein-coding domains from all 43 proteomes considered in this analysis (core CCs), 160,431 CCs exclusively included protein-coding domains from a single proteome (accessory CCs), and 347,551 CCs corresponded to the pan proteome of the dinoflagellates included in the analysis (Fig. 1*A*). We extrapolated the trend of the core proteome CC number using a non-linear regression model. The best-fit function was *y* = *a* / *x*, with *y* the predicted number of core CCs, *x* the number of proteomes and *a* an estimated parameter. For 2 to 43 proteomes, this model had a correlation of 0.97 to our data (p-value of estimated parameter *a* < 2e-16). The extrapolation of the number of core CCs for 50, 60 and 70 proteomes were 170, 144 and 123 CCs respectively, without displaying a saturation to a fixed number of core CCs. The Pielou diversity indices calculated to explore CC composition had a mean value of 0.96, indicating not only that core CCs were composed of all proteomes by definition, but that they were also evenly structured, rarely being dominated by a single proteome.

**Fig. 1:**
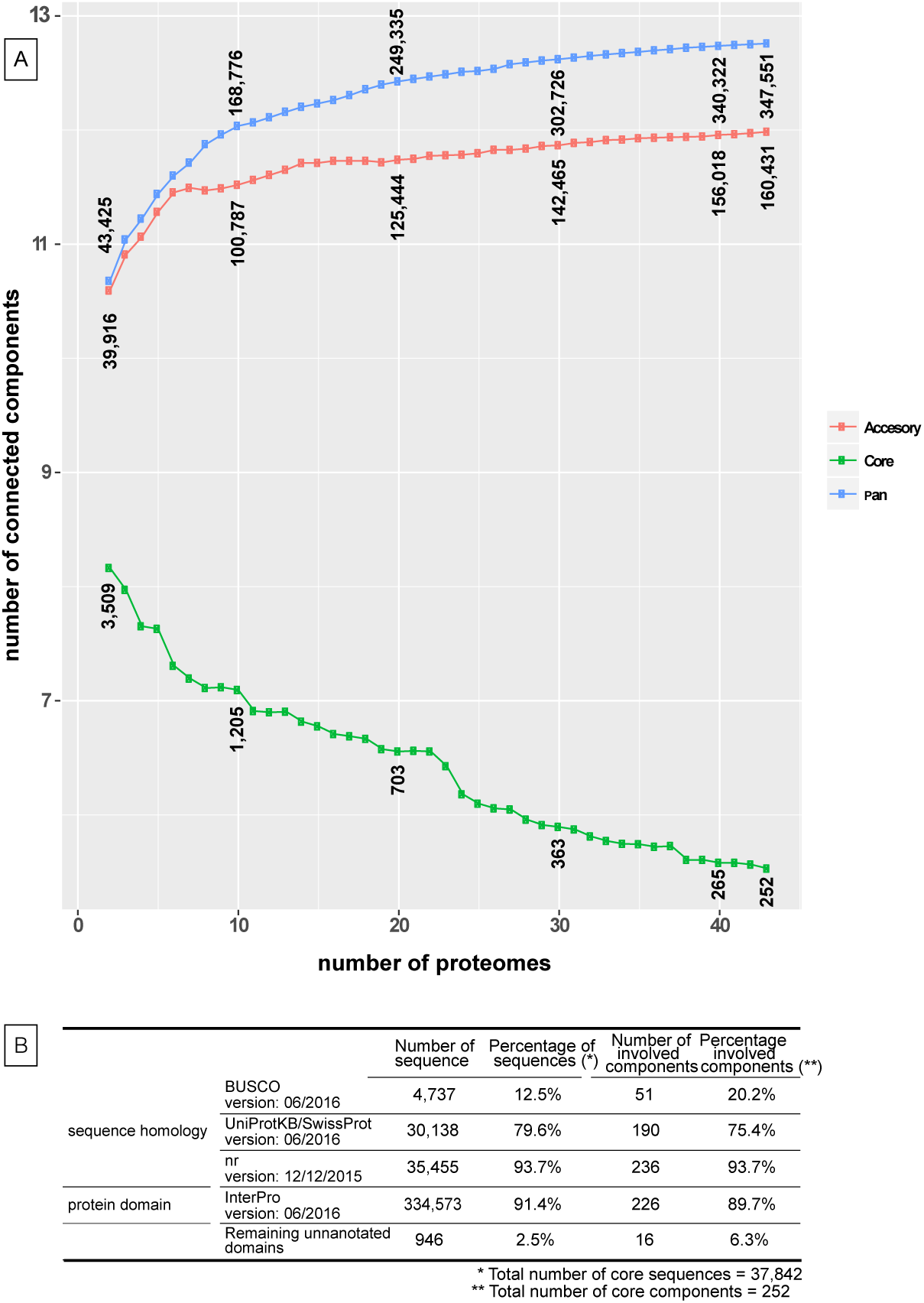
(A) Number of connected components (CCs) in the core (green), accessory (red) and pan (blue) dinoflagellate proteomes, considering 2 to 43 proteomes. (B) Comparison of the 37,842 protein domains included in the 252 core dinoflagellates CCs to BUSCO, UniProtKB/Swiss-Prot and nr databases. The number and percentage of core sequences with at least one match in each database, and the number and percentage of their corresponding CCs.

Functional annotation using InterPro revealed that 91,4% of core protein-coding domains matched to the InterPro database. According to GOslim functional annotations, core CCs had an important contribution of protein-coding domains annotated as “ribosomal proteins” having a role in RNA translation (*i.e*. 7,968 of 37,842 core protein-coding domains) (Fig. S2). Other main functional annotations occurring in core CCs were protein phosphorylation (1,752 protein-coding domains), proteins involved in signal transduction (1,133 protein-coding domains) and cell redox homeostasis (562 protein-coding domains), the rest being composed of a variety of functions represented by few protein-coding domains (Fig. S2). The 37,842 protein-coding domains belonging to the 252 core CCs were further analyzed by comparison to other reference databases to identify protein-coding domains that are shared among the 43 dinoflagellate proteomes. The proportion of matching protein-coding domains reached up to 12.5% (involved in 51 CCs), 79.6% (involved in 190 CCs) and 93.7% (involved in 236 CCs) against BUSCO (Simão *et al.* 2015), UniProtKB/Swiss-Prot and nr, respectively (Fig. 1*B*). A total of 16 CCs (*i.e.* 946 protein-coding domains) from the core proteome did not have a match in any of the databases explored (Fig. 1*B*) (Tab. S2). The 101 orthologous alignments used for a recent phylogeny of dinoflagellates (Janouškovec *et al.* 2016) were compared to the protein-coding domains from the 252 core CCs. Results show that 1606 protein-coding domains from 46 CCs matched with at least one of the 101 orthologous alignments (Fig. S3), and that domains from our 16 unknown core CCs were not included in these 46 CCs.

### Functional trait investigations

In the SSN based on the 46 proteomes, the number of CCs exclusively composed of protein-coding domains from species tagged for a functional trait (*i.e.* trait-CCs) has been reported for each trait investigated (Tab. S3-S12), as well as the percentage of trait-CCs that are annotated (*e.g.* an annotated trait-CC includes at least one functionally annotated protein-coding domain). As expected considering the taxonomic coverage of our dataset, the analysis revealed a maximum number of trait-CCs for the “chloroplast” trait (336,099 CCs) and a minimum for the “parasitism” trait (826 CCs). The “chloroplast” trait had the highest percentage of annotated trait-CCs (93%) while the “parasitism” trait had the lowest (23%) (Fig. S4). Among the trait-CCs, a total of 5 “harmful for human” trait-CCs regrouping protein-coding domains of 7 of 14 possible proteomes were detected. Likewise, we found 2 “symbiosis” trait-CCs including 8 of 12 possible proteomes (Tab. S4 & S6).

### Identification of toxin related sequences in “harmful for human” trait-CCs

Well-described proteins involved in dinoflagellate toxicity, the polyketide synthases (PKS) and saxitoxins (STX) were sought within the “harmful for human” trait-CCs. 36 protein-coding domains homologous to PKS were identified in 17 “harmful for human” trait-CCs (composed of a total of 45 protein-coding domains) (Tab. S13). On the other hand, 646 protein-coding domains homologous to PKS were found in 165 CCs (composed of a total of 1,144 protein-coding domains) belonging to the contrasting non-“harmful for human” trait-CCs. All protein-coding domains from trait-CCs in which PKS homologues were detected (*i.e.* 45 + 1,144 = 1189 protein coding domains) had either a Thiolase-like functional annotation (1,159 protein-coding domains), which corresponds to the superfamily of KS enzyme domains of PKS, or lacked annotation (30 protein-coding domains) according to the InterPro database. The *sxtA* and *sxtG* genes have been reported to be involved in the STX biosynthesis pathway. Based on 117 *sxtA1-4* and 20 *sxtG* reference gene sequences, no target protein-coding domain matched to “harmful for human” trait-CCs (Tab. S14). In contrast, 99 and 3 unique protein-coding domains were identified for *sxtA1-4* and *sxtG* respectively, in non-“harmful for human” trait-CCs. *sxtA1-4* hits involved protein-coding domains from 20 CCs (composed of 166 protein-coding domains), and *sxtG* hits belonged to 1 CC composed of 3 protein-coding domains. Of the 166 *sxtA1-4* homologues, 156 protein-coding domains had InterPro annotations related to *sxtA* domains (*i.e.* pyridoxal phosphate-dependent transferase, PKS, GNAT domains, Acyl-CoA N-acyltransferase) and the remaining 10 protein-coding domains were unannotated. A single InterPro functional annotation was found for the CC involving *sxtG* homologues (of the 3 protein-coding domains forming the CC) and corresponded to an amidinotransferase domain that is known as a *sxtG* protein domain (Tab. S14).

We investigated the GO functional annotations of “harmful for human” trait-CCs. At the cellular component functional level, “membrane” and “integral component of membrane”, protein-coding domains represented 51% (3017 out of 5,998) and 27% (1,646 out of 5,998) of annotated protein-coding domains, respectively (Fig. 2*A*). At the biological process annotation level, 14% (1,672 out of 11,187) of protein-coding domains were linked to “ion transport” (Fig. 2*A*). At the molecular function annotation level, 24% (5,151 out of 21,337) corresponded to “protein binding” protein-coding domains (Fig. 2*A*). Differential composition of functional annotations between proteomes of “harmful for human” compared to non-“harmful for human” species was investigated to unveil functions that are more likely to be observed in the proteome of toxic species. The pair differences of each function occurring in “harmful for human” and non-“harmful for human” trait-CCs showed that “ion transport” protein domains occured 7 times more often in “harmful for human” trait-CCs. We also noticed that pentatricopeptide repeat, C2 domain, P-loop containing nucleoside triphosphate hydrolase, Pyrrolo-quinoline quinone beta-propeller repeat, Quinonprotein alcohol dehydrogenase-like and Thrombospondin type 1 repeat domains occurred from 1 to 2 times more often in “harmful for human” trait-CCs (Fig. 2*B*). The core “harmful for human” trait-CCs (*i.e*. CCs that are composed of protein-coding domains from most toxic species representatives) was investigated to reveal functions that are shared among toxic species only (Fig. 2*C*). We identified 5 of such core “harmful for human” trait-CCs, corresponding to a total of 49 protein-coding domains. These core trait-CCs encompassed 7 of 14 toxic dinoflagellate proteomes considered in our analysis. Not a single of these 49 protein-coding domains had a GO annotation. Based on InterPro functional annotations, 3 of the 5 CCs are respectively composed of 14 “nucleotide-binding alpha-beta plait” protein-coding domains, 7 “P-loop containing nucleoside triphosphate hydrolase” protein-coding domains and 8 “nucleotide-diphospho-sugar transferase” protein-coding domains. Two of these 5 CCs were entirely composed of 7 and 15 unannotated protein-coding domains. The taxonomic distribution of the 49 protein-coding domains is represented by: 30 protein-coding domains from the *Alexandrium* genus (61%), 2 from *Dinophysis acuminata*, 4 from *Protoceratium reticulatum*, 6 from *Gambierdiscus australes*, 4 from *Pyrodinium bahamense* and 3 from *Lingulodinium polyedra*.

**Fig. 2:**
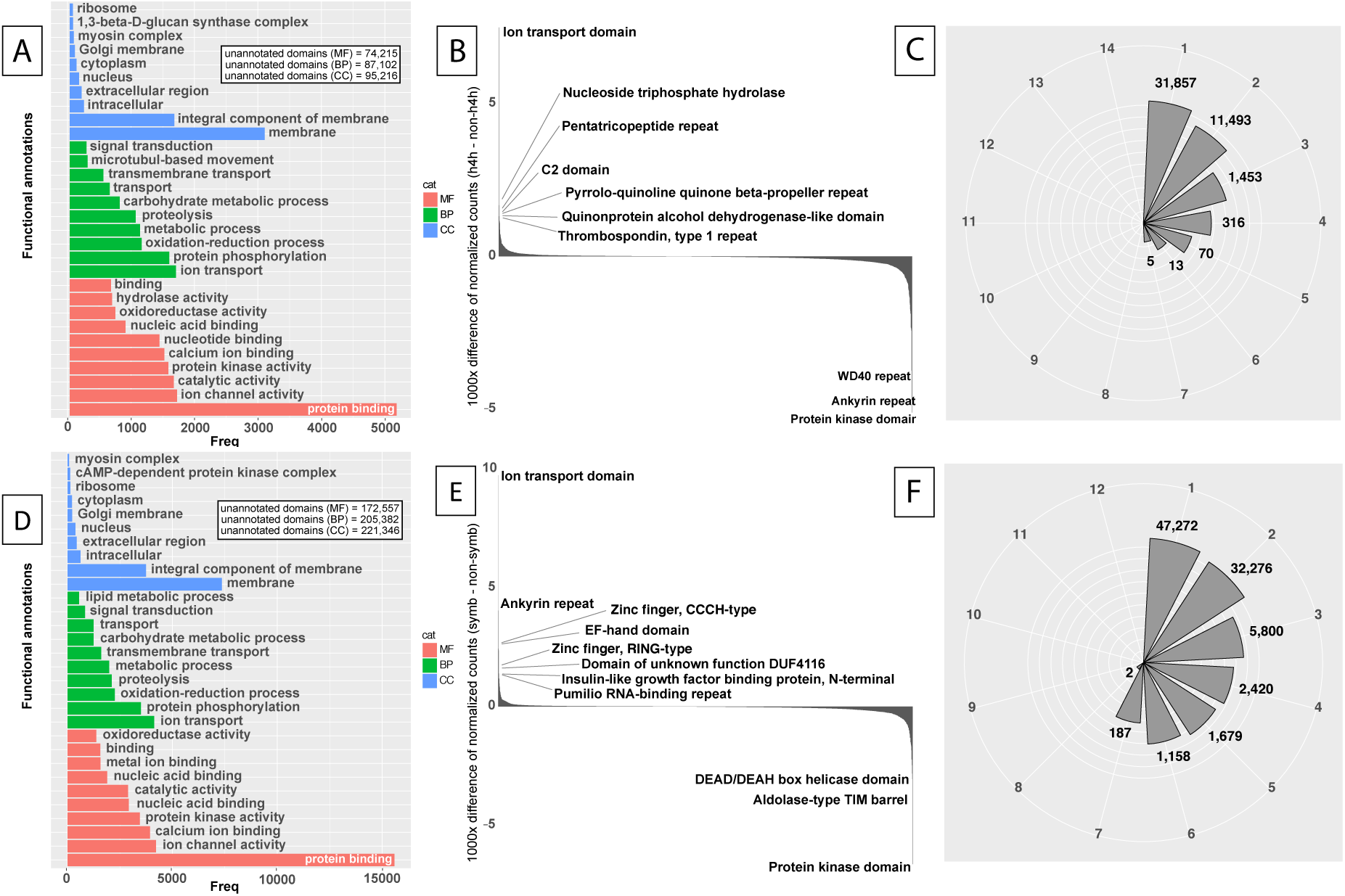
(A and D) Top 10 functional annotations (GOslim levels) of sequences belonging to the 45,207 “harmful for human” trait-CCs (A) and to the 90,794 “symbiosis” trait-CCs (D). (B and E) Differential composition of functional annotations between “harmful for human” and non-“harmful for human” trait-CCs (B) and “symbiosis” and non-“symbiosis” trait-CCs (E). (C and F) The circular barplot shows the number of connected components that include 1 to 14 proteome(s) of the transcriptomes assigned to toxic species (C) and the number of connected components that include 1 to 12 proteome(s) of the transcriptomes assigned to symbiotic species (F).

31,496 “harmful for human” trait-CCs, composed in total of 70,359 protein-coding domains, completely lacked functional annotations according to Interproscan. Alignments (with an e-value of 1e-3) to nr database revealed 6,103 hits including 283 protein-coding domains with sequence identity higher than 80%. These 283 protein-coding domains originated from 157 CCs which were composed of 359 protein-coding domains in total. Finally, we identified functions for 157 “harmful for human” trait-CCs while 25,393 “harmful for human” trait-CCs remained without functional annotation.

### Exploration of “symbiosis” trait-CCs

A large range of dinoflagellates, including symbiotic and non-symbiotic species, express genes identified in the literature as potentially involved in symbiotic processes (Tab. S15). 150 of these gene sequences were compared to protein-coding domains included in “symbiosis” trait-CCs. Alignments revealed 8 protein-coding domains from 5 “symbiosis” trait-CCs. These 8 protein-coding domains were identified as belonging to proteins involved in symbiosis establishment (nodulation protein noIO and phosphoadenosine phosphosulfate reductase), cell recognition processes (merozoite surface protein), and highlighted in cnidarian-algal symbiosis (peroxiredoxin, ferritin) (Tab. S15). Similarly, 71 protein-coding domains (spread across 21 CCs) matching symbiosis related query sequences were found in non-“symbiosis” trait-CCs. Functions of these 71 protein-coding domains are involved in symbiosis establishment (P-type H+-ATPase, phosphoadenosine phosphosulfate reductase), cell recognition processes (merozoite surface protein 1) and exposed in cnidarian-algal symbiosis (superoxide dismutase, catalase, peroxiredoxin, glutathione peroxidase, g-glutamylcysteine synthetase).

We explored GO functional annotations from all “symbiosis” trait-CCs (Fig. 2*D*). At the cellular component level, 83% of annotated protein-coding domains (11,103 out of 13,298) were “membrane” protein-coding domains. At the biological process level, 21% (4,131 out of 19,380) were “ion transport” protein-coding domains and 18% were involved in “protein phosphorylation”. At the molecular function level, 39% (15,585 out of 39,293) were annotated as “protein-binding” protein-coding domains, 10% and 9.9% were involved in “ion channel activity” and “calcium ion binding”, respectively. Pair differences of normalized counts for each possible function that appeared in proteomes of both symbiotic and non-symbiotic species revealed 4 annotations that occured 2 to 10 times more within “symbiosis” trait-CCs. These were protein-coding domains containing an ion transport domain, ankyrin repeat domains, EF-hand domain and zinc finger, and CCCH-type annotations were 9.7, 4, 2.4, and 2.3 times more numerous, respectively, in “symbiosis” trait-CCs (Fig. 2*E*).

Our attempt to identify core CCs of the “symbiosis” trait yielded 2 “symbiosis” trait-CCs involving a maximum of 8 distinct proteomes of symbiotic species and 187 “symbiosis” trait-CCs involving 7 proteomes (of the 12 proteomes available for symbiotic species) (Fig. 2*F*). GO annotations of these 189 core “symbiosis” trait-CCs revealed that the major part of the protein-coding domains (*i.e.* 1400 out of 1896) could not be functionally annotated. Among those that could be annotated, 73.8% of the protein-coding domains corresponded to “membrane proteins” (cellular component). The remainder corresponded to “proteins of photosystem I”, “extracellular region” and “spliceosomal complex”. With respect to biological process, 31.9% were involved in ion transport while 23.8% were involved in proteolytic processes (Tab. S16). Looking at taxonomic distribution, we found that the 189 CCs were dominated by protein-coding domains belonging to *Symbiodinium* spp. (1679 protein-coding domains from the 187 CCs and 13 protein-coding domains from the 2 CCs). *Pelagodinium beii* was represented by 211 protein-coding domains spread across the 187 CCs. Finally, few protein-coding domains (respectively 8 and 1) of *Gymnodinium radiolariae* and *Brandtodinium nutricula* were found within the 189 “symbiosis” trait-CCs.

The 52,491 “symbiosis” trait-CCs (composed of 130,673 protein-coding domains that lack InterPro functional annotation) were compared to nr database alignments (with an e-value of 1e-3) revealing 495 protein-coding domains with sequence identity higher than 80%. These 495 protein-coding domains belonged to 298 CCs, which were composed of 919 protein-coding domains in total. Finally, 52,193 “symbiosis” trait-CCs were completely unannotated.

## DISCUSSION

### Pipeline efficiency & sequence similarity network

Our *de novo* assembly and downstream pipeline analysis of multiple dinoflagellate transcriptomes overcame several biases inherent to *de novo* assembly processes (Fig. S5). For instance, the protein-coding domain prediction step we performed contributed to selection of transcripts in which ORFs and protein domains were detected but also allowed removal of truncated or chimeric transcripts (Yang & Smith 2013). Protein-coding domains derived from high quality transcriptomes enabled construction of sequence similarity networks to focus on shared protein-coding domains among multiple proteomes. Considering our 46 proteomes, a mean value of 60,661 protein-coding domains was found, which is consistent with the previously estimated range of 34,156 to 75,461 protein-coding genes in dinoflagellates (Murray *et al.* 2016). The median length of the protein-coding domains was 307 bp, also consistent with the median protein length of 361 bp reported based on genomes of 5 model eukaryote species (*Homo sapiens*, *Drosophila melanogaster*, *Caenorhabditis elegans*, *Saccharomyces cerevisiae* and *Arabidopsis thaliana*) (Brocchieri & Karlin 2005).

Sequence similarity networks represent an informative and pragmatic way to study massive datasets (Atkinson *et al.* 2009; Alvarez-Ponce *et al.* 2013; Cheng *et al.* 2014; Forster *et al.* 2015; Méheust *et al.* 2016). In (Cheng *et al.* 2014), 84 genome-derived proteomes of prokaryotes (*i.e.* 128,628 sequences) were used to study the impact of redox state changes on their gene content and evolution. The authors found that the core CCs revealed a correlation between their network structure and differences in respiratory phenotypes. Our SSN has allowed simultaneous exploration of 46 transcriptome-derived proteomes (1,275,911 sequences), including their overwhelming “dark matter” (*i.e.* here protein-coding domains totally lacking functional annotation). High identity and coverage threshold values used in our analyses to filter alignments ensured that only high quality alignments were included in the network (Bittner *et al.* 2010). The integration of 4 new dinoflagellate proteomes represented an increase of 14% of protein-coding domains in the SSN and overall the dataset represents the most comprehensive picture to date of the genomic potential of dinoflagellates. This new resource and comparative genomic approach allow generation and testing of original hypotheses about the genomic basis for evolutionary history and life style, functional traits, and specificities of dinoflagellates.

### Large-scale comparison of dinoflagellate proteomes

The SSN analyses allowed characterization of the core and accessory proteomes for this large dataset of non-model organisms. Because our analysis relied on a *de novo* assembled, transcriptome-derived, proteome SSN rather than classical knowledge-based genomics, it also promoted discovery of new CCs, each of which can be functionally assimilated to a single putative conserved protein-domain (pCD) in such non-model organisms (Lopez *et al.* 2015) (Fig. S6).

The core dinoflagellate proteome identified in our analysis was composed of 252 pCDs (Fig. 1*A*), a size that falls in the range of the latest estimates for bacteria (352 core genes) (Yang *et al.* 2015) and eukaryotes (258 core genes in CEGMA, and more recently 429 single-copy orthologs in BUSCO) (Parra *et al.* 2007; Simão *et al.* 2015). The extrapolation of the number of core CCs does not saturate, suggesting that the number of core CCs for dinoflagellates could be less than 256. It also suggests that dinoflagellates expressed only a fraction of core eukaryote genes referenced in databases. Our comparative analysis with the most up-to-date eukaryotic orthologous gene database BUSCO strongly stresses the need to generate more gene and protein data for non-model marine organisms in order to populate reference databases (Armengaud *et al.* 2014). The small overlap between core dinoflagellate pCDs identified here and the BUSCO database suggests that essential functions expressed by dinoflagellates are distantly related to those of current model eukaryotes.

Our SSN constitutes a strong basis for exploration and refinement of functional annotations as our dataset encompassed a broad range of dinoflagellate taxa according to recent phylogenetic analyses (Bachvaroff *et al.* 2014; Janouškovec *et al.* 2016). However, the identified core proteome can only be considered partial as our dataset *i-* did not include representatives of all described dinoflagellate lineages, and *ii-* relied on transcriptomic (*i.e.* gene expression) data that can vary according to eco-physiological conditions and/or life-cycle stage. The content of our SSN can be updated permanently to refine these estimates as new dinoflagellate genomic data are accumulated (Shoguchi *et al.* 2013; Lin *et al.* 2015; Aranda *et al.* 2016). 236 (93%) core CCs involving one or more functionally annotated protein-coding domains (Fig. 2*B*) can be exploited to extend annotation to other aligned protein-coding domains within each CC, therefore leading to a more comprehensive pCD description. For instance, looking for the HSP70 conserved protein domain, which is ubiquitous in all eukaryotic organisms (Germot & Philippe 1999), we found 320 protein-coding domain sequences annotated as HSP70, all belonging to a single CC composed of 328 protein-coding domain sequences. The 8 remaining protein-coding domain sequences were either imprecisely annotated as chaperone DnaK (1 sequence), cyclic nucleotide-binding domain (2 sequences), heat shock protein 70 family (3 sequences) or annotation was simply missing (2 sequences) (Tab. S17). As this CC was 97% represented by HSP70 annotation, it is reasonable to extend this annotation to all protein-coding domain sequences forming the connected component. Considering only CCs that were at least half composed of annotated protein-coding domain sequences, this approach could be applied to complement the functional characterization of 49 CCs (583 unannotated protein-coding domains) through extension of functional annotations.

Relatively few previous studies have relied on functional aspects such as protein alignments to investigate dinoflagellate phylogeny. A recent study used for the first time a multi-protein dataset providing a robust phylogeny for dinoflagellates (Janouškovec *et al.* 2016). The comparison of the 101 orthologous alignments used in (Janouškovec *et al.* 2016) with our 252 pCDs revealed 206 additional core pCDs that are good candidates for refining dinoflagellate phylogeny, increasing by nearly 200% the quantity of information available for such studies.

Among the 176,958 distinct CCs entirely composed of unannotated protein-coding domains in the total network, 16 CCs or pCDs (composed of 946 protein-coding domain sequences) belonged to our core dinoflagellate proteome (Fig. 1*B*). This highlights the fact that many fundamental genomic features remain to be characterized in this group. These unknown groups of homologous domains are excellent potential candidate markers to further investigate dinoflagellate genomics at a broad scale and might also be useful for identification of dinoflagellates within complex environmental genomic datasets.

### Investigating harmful dinoflagellates

Toxic dinoflagellates represent about 80% of toxic eukaryotic phytoplankton species (Janouškovec *et al.* 2016). Production of toxins by dinoflagellates is well known and can cause major health and economic problems. *Karenia brevis*, for example, is known to produce brevetoxins which cause fish mortality and can affect human health through the consumption of contaminated seafood or direct exposure to harmful algal blooms (HABs) (Flewelling *et al.* 2005). To date, several dinoflagellate toxins have been chemically and genetically characterized (Wang 2008; Kellmann *et al.* 2010; Stüken *et al.* 2011; Cusick & Sayler 2013). In our SSN analyses, protein-coding domains homologous to PKS were identified in CCs composed of domains from both “harmful for human” and non-“harmful for human” species. This result validates a previous report that PKS proteins are not exclusive to toxic species (Kohli *et al.* 2016). PKS are in fact involved in the production of a variety of natural products such as small acids, acetyl-CoA or propionyl-Co (Khosla *et al.* 2014). Spreading information among unannotated protein-coding domains in both “harmful for human” and non-“harmful for human” trait-CCs in which PKS were identified allowed extension of the potential PKS-like annotation to 9 protein-coding domains from “harmful for human” trait-CCs and 498 protein-coding domains from non-“harmful for human” trait-CCs. PKS protein-coding domains for 4 extra species (*Alexandrium catenella*, *Kryptoperidinium foliaceum*, *Protoceratium reticulatum* and *Crypthecodinium cohnii*) were also detected compared to the database from (Kohli *et al.* 2016) (Tab. S13) confirming that synthesis of PKS proteins is not exclusive to toxic species (Kohli *et al.* 2016).

With respect to saxitoxin production, we did not detect either *sxtA* or *sxtG* related protein-coding domains in “harmful for human” trait-CCs which suggests that such proteins are not exclusively expressed by toxic species. Robust alignments for both *sxtA* and *sxtG* protein-coding domains in non-“harmful for human” trait-CCs were found. Our results differed somewhat from those of a previous study (Murray *et al.* 2015) based on the same initial dataset from the MMETSP. Specifically, we were not able to detect *sxtA* in 7 species in which this gene was detected in (Murray *et al.* 2015) and in contrast *sxtA* domains were detected in 9 species in which the protein was not detected previously (Murray *et al.* 2015) (Tab. S15). These differences may be due to the use of distinct *de novo* assembly tools and pCD prediction processes, illustrating the requirement to ultimately combine *in vitro* and *in silico* methods in order to unambiguously characterize toxic species. Nevertheless, our results seem consistent with biological knowledge since the expression of both *sxtA* and *sxtG* proteins, which are involved in toxin biosynthesis process in toxic species (Hackett *et al.* 2013), has already been revealed for *P. bahamense* and *G. catenatum* that are known to be STX-producing dinoflagellates and reported as toxic species (Tab. S14). We also confidently detected 2 protein-coding domains including *sxtA* domains and 1 protein-coding domain including *sxtG* domains in *P. beii*, an *a priori* non-toxic symbiotic species that has never been reported as a STX-producer. This is consistent with the fact that *sxtG* has previously been identified in non-toxic species (Orr *et al.* 2013). The detailed investigation of the 2 first domains sharing similarity to *sxtA* showed that one shares an aminotransferase domain with *sxtA* (Hackett *et al.* 2013) and the second shares GNAT domains with *sxtA*. These results alone are not sufficient to prove toxin production and toxicity tests must be performed *in vitro* to confirm the synthesis of saxitoxins by *P. beii*. Previous studies showed that two forms of *sxtA* (long and short) are present in some *Alexandrium* genera (Stüken *et al.* 2011) and that the shorter form is not related to toxin production (Murray *et al.* 2015). We were not able to associate the domains identified in our study to either the long or short form of *sxtA*, and we cannot exclude the possibility that the *sxtA* domain identified could belong to a molecule that is synthesized through the saxitoxin biosynthesis pathway but that is not a functional saxitoxin. From an evolutionary point of view, as PKS and STX genes are also found in species currently described as non-toxic, it seems that like for snake venoms, dinoflagellate toxins evolved by recruitment of genes encoding regular proteins followed by gene duplication and neo-functionalization of the domains (Vonk *et al.* 2013).

Based on exploration of the SSN for putative coding domains specific for “harmful for human” trait-CCs, we found that membrane located protein domains and more specifically ion transport protein domains were important components characterizing toxic species. This is in agreement with reports that ion channel proteins and proteins involved in neurotransmission are mediators of dinoflagellate toxicity (Wang 2008; Cusick & Sayler 2013). For our SSN analysis, we made the assumption that the more species are represented in a CC, the more the corresponding gene set is likely to be specific for this particular functional trait. In the case of “harmful for human” trait-CCs we noticed that 2 of the 5 CCs with the most toxic representatives (*i.e.* 7 species) were exclusively composed of unannotated domains, representing essential functions constitutively expressed by toxic species only and for which further investigations are required to better characterize toxic dinoflagellates.

### Focus on symbiosis

The “symbiotic” gene set compiled from the literature based on their involvement in the establishment and maintenance of symbiosis (Lehnert *et al.* 2014; Lin *et al.* 2015) was found here in both “symbiosis” trait-CCs and in non-“symbiosis” trait-CCs (Tab. S15), suggesting that these proteins are constitutively expressed by all dinoflagellate species. This result may reflect the fact that the transcriptomes of dinoflagellate strains were not directly isolated from symbiotic conditions, but rather from their free-living stages maintained in culture. Symbiotic genes identified from the literature were originally inferred from studies on holobionts (*i.e.* host and symbionts), but proved here not to be exclusive to symbiotic dinoflagellates when performing global comparison of multiple datasets.

Functional annotations of “symbiosis” trait-CCs revealed an overall clear domination of proteins involved in phosphorylation and ion transport domains (*e.g.* sodium, potassium and calcium ion channel proteins) located within membrane compartments (Fig. 2*D*). The 4 most prominent functions that occured 2 to 10 times more often in “symbiosis” trait-CCs than in non-“symbiosis” trait-CCs (Fig. 2*E*) were related to ion transport domains and regulation processes. The results indicate that the two functions are constitutively more expressed in symbiotic compared to non-symbiotic species. Protein phosphorylation is known to take part in cellular mechanisms in response to the environment (Day *et al.* 2016) and play a key role in signal transduction to other cells in plant parasitism and symbiosis models (Lionetti & Metraux 2015). The specific dominant presence of ion transport domains (also involved in cell signaling and cell adaptation to the environment) in symbiotic dinoflagellates could represent a constitutive characteristic of symbiotic species facilitating establishment and maintenance of the symbiosis. Notably, the role of ion channel proteins has been highlighted as essential in plant root endosymbiosis (Charpentier *et al.* 2008; Matzke *et al.* 2009). This suggests that symbiotic species are likely to be constitutively better equipped for environmental adaptations.

Exploring the SSN, we found that 45% of the overall protein-coding domains associated to symbiotic species were functionally annotated (Tab. S19) which implies that a large suite of uncharacterized functions are specifically associated to symbiotic dinoflagellates. 129,754 protein coding domains from 52,193 “symbiosis” trait-CCs remained unannotated according to the InterPro and nr databases. We found 187 “symbiosis” trait-CCs composed of 7 of 12 possible symbiotic species and 2 CCs composed of 8 of the 12 (Fig. 2*F*). Protein-coding domains from 2 of the 3 newly added symbiotic proteomes were found in both the 187 and 2 pCDs, contributing to revealing pCDs specific of symbiotic species. The 2 “symbiosis” trait-CCs encompassing the 8 distinct species were exclusively composed of unannotated domains, suggesting that these 2 CCs represent pCDs with fundamental, yet unknown, functions constitutively expressed by symbiotic species. Overall, our analyses demonstrate that SSN has significant potential to reveal the variety of annotated and unknown pCDs that constitute good candidates for further study to characterize and understand the genomic basis of symbioses involving dinoflagellates.

## M&M

### Dataset building

The dataset used in our study included all dinoflagellate transcriptomes available in the MMETSP project repository (http://marinemicroeukaryotes.org/resources) as well as 4 transcriptomes generated for this study (more details in the following section) (Fig. S7). This dataset represented transcriptomes of 47 distinct species from 35 genera, 19 families, and 11 of the 21 current dinoflagellate taxonomic orders according to the taxonomic framework of the WoRMS database (http://www.marinespecies.org/index.php) (Tab. 1). Taxonomy and functional trait information (i.e. chloroplast occurrence and origin, trophic mode, harmfulness for humans, ability to live in symbiosis, to perform kleptoplasty, to be a parasite or to be toxic for fauna) were indicated for each organism considered (Tab. 1).

### Culturing and RNA sequencing for four dinoflagellate strains

Free-living clonal strains of the dinoflagellate species *Brandtodinium nutricula* (RCC3468) (Probert *et al.* 2014) and *Gymnoxanthella radiolariae* (RCC3507) (Yuasa *et al.* 2016) isolated from symbiotic Radiolaria, *Pelagodinium beii* (RCC1491) (Siano *et al.* 2010) isolated from a foraminiferan host, and the non-symbiotic *Heterocapsa* sp. (RCC1516) were obtained from the Roscoff Culture Collection (www.roscoff-culture-collection.org). Triplicate 2-L acid-washed, autoclaved polycarbonate Nalgene bottles were filled with 0.2 micron filter-sterilized (Stericup-GP, Millipore) seawater with K/2 (-Tris,-Si) medium supplements (Keller *et al.* 1987) and inoculated with an exponentially growing culture of each strain. All cultures were maintained at 18°C, ~80 µmol photon m^−2^ s^−1^ light intensity and 14:10 light:dark cycle. Cell abundance was monitored daily by flow cytometry with a FACSAria flow cytometer (Becton Dickinson, San José, CA, USA) and derived cell division rates were used to monitor the growth phase of the culture. Light and dark phase samples for transcriptome analyses were taken from exponential and stationary phase cultures. 100 mL aliquots from each culture were filtered onto 3 micron pore-size polycarbonate filters with an autoclaved 47 mm glass vacuum filter system (Millipore) and a hand-operated PVC vacuum pump with gauge to maintain the vacuum pressure below 5 mm Hg during filtration. The filter was then placed in a sterile 15 mL falcon tube filled with ca. 5 ml TriZol and stored at −80°C.

Total RNA was purified directly from the filters stored in TriZol using the Direct-zol RNA Miniprep kit (ZymoResearch, Irvine, CA). First, the tube containing the filter immersed in TriZol was incubated for 10 min at 65°C. Then, after addition of an equal volume of 100% EtOH and vortexing, the mixture was loaded into a Zymo-SpinIIC column and centrifuged for 1 min at 12,000 *g*. The loading and centrifugation steps were repeated until exhaustion of the mixture. RNA purification was completed by prewash and wash steps following the manufacturer's instructions and RNA was directly eluted in 45 µL nuclease-free water. The in-column DNAse step was replaced by a more efficient post-extraction DNAse treatment using the Turbo DNA-free kit (Thermo Fisher Scientific, Waltham, MA) according to the manufacturer's rigorous DNase treatment procedure. After two rounds of 30 minutes incubation at 37°C, the reaction mixture was purified with the RNA Clean and Concentrator-5 kit (ZymoResearch) following the procedure described for retention of >17nt RNA fragments. Total RNA, eluted in 20 µl nuclease-free water, was quantified with RNA-specific fluorimetric quantification on a Qubit 2.0 Fluorometer using Qubit RNA HS Assay (ThermoFisher Scientific). RNA quality was assessed by capillary electrophoresis on an Agilent Bioanalyzer using the RNA 6000 Pico LabChip kit (Agilent Technologies, Santa Clara, CA).

RNA-Seq library preparations were carried out from 1 µg total RNA using the TruSeq Stranded mRNA kit (Illumina, San Diego, CA), which allows mRNA strand orientation. Briefly, poly(A)^+^ RNA was selected with oligo(dT) beads, chemically fragmented and converted into single-stranded cDNA using random hexamer priming. Then, the second strand was generated to create double-stranded cDNA. Strand specificity was achieved by quenching the second strand during final amplification thanks to incorporation of dUTP instead of dTTP during second strand synthesis. Then, ready-to-sequence Illumina libraries were quantified by qPCR using the KAPA Library Quantification Kit for Illumina libraries (KapaBiosystems, Wilmington, MA), and library profiles evaluated with an Agilent 2100 Bioanalyzer (Agilent Technologies). Each library was sequenced using 101 bp paired-end read chemistry on a HiSeq2000 Illumina sequencer.

### Data filtering and *de novo* assembly

For each strain considered (Tab. S7 and our 4 strains), sequenced reads were pooled resulting in 60 datasets, then filtered using Trimmomatic (Tab. 1) (34). Reads with quality below 30 Q on a sliding window size of 10 were excluded. Remaining reads were assembled with the *de novo* assembler Trinity version 2.1.1 (Grabherr *et al.* 2011) using default parameters for the paired reads method. Of the initial 60 transcriptome datasets (56 from the MMETSP repository and 4 produced in this study), 57 were successfully assembled. The assembly process could not be completed properly for 3 datasets due to incompatibility between the version of the assembly software and the datasets (*Karenia brevis* strain CCMP 2229, Wilson SP1 and SP3 as a combined assembly, *Oxyrrhis marina* strain CCMP1795 and *Symbiodinium kawaguti* strain CCMP2468) (Tab. 1).

Assembled transcripts were evaluated based on: (i) sequence metrics, and (ii) read remapping rates calculated respectively with homemade scripts and Bowtie 2 in local mode (Langmead *et al.* 2009) (Tab. 1). Based on these assembly analyses, two classes of assembly quality were defined: those with >30,000 transcripts with a N50 > 400 bp and read remapping rate >50% were tagged as “high quality” transcriptomes whereas the rest were tagged as “low quality” transcriptomes. An exception was made for one poor quality transcriptome corresponding to the species *Oxyrrhis marina* (LB1974 and NA strain) composed of 18,275 assembled transcripts that was intentionally tagged as a “high quality” transcriptome because this basal species holds a key evolutionary and ecological position among dinoflagellates (Montagnes *et al.* 2011; Lee *et al.* 2014; Bachvaroff *et al.* 2014).

### Coding domain prediction and functional annotation

For each transcriptome, coding domain prediction of assembled transcripts was conducted with Transdecoder version 2.0.1 (Haas *et al.* 2013) to obtain peptide sequences of corresponding domains. We defined each set of predicted protein domains as a proteome. The optional step of Transdecoder consisting in the identification of ORFs in the protein domain database Pfam was not executed in order to avoid a comparative approach that would result in a limited discovery of new sequences. The predicted coding domains were then processed with the Interproscan 5 functional annotation program version 5.11-51.0 (Jones *et al.* 2014) to scan for protein signatures. Default parameters were used to obtain each proteome. Finally, to get a broad overview of the ontology content of our datasets, GO slims were retrieved from the Gene Ontology Consortium to build a summary of the GO annotations without the detail of the specific fine-grained terms (http://geneontology.org/page/go-slim-and-subset-guide).

### Sequence similarity network

A sequence similarity network (SSN) is a graph in which vertices are genomic sequences and the edges represent similarity between sequences. A SSN is composed of connected components (CC) (subgraphs or subnetworks, including at least two vertices disconnected from other subgraphs in the total network). As information can be linked to sequences (e.g. in our study: taxonomy, functional annotation, functional traits), the SSN and its structure can be explored accordingly. Using predicted protein domain sequences, a SSN was constructed with the BLASTp alignment method (Altschul *et al.* 1990) with an e-value of 1e-25 using the DIAMOND software (Buchfink *et al.* 2015). Similarities satisfying query and subject sequence coverages higher than 80% were kept.

Whenever predicted coding domains aligned together forming a CC it can be assumed that they potentially share a similar molecular function (Marchler-Bauer *et al.* 2005) and form putative conserved domains (pCDs). SSN exploration and analyses were performed using personal scripts and functions implemented in the igraph R package (Csárdi & Nepusz 2006). Diverse biological information related to the species considered were mapped on each vertex, and missing information were marked as <NA>.

In our approach, CC number, structure and composition were impacted when edge sequence identity cut off was shifted. We thus tested different similarity thresholds and chose an optimal threshold according to the two following criteria: maximizing the number of large CCs (i.e. minimum of 30 vertices) and the number of CCs involving a single homogeneous functional annotation (i.e. a unique GOslim term at the Biological Process level). An optimal sequence identity threshold at 60% similarity with our dataset was inferred (Fig. S1). As a last filtering step, we chose to consider only vertices of proteomes derived from “high quality” tagged transcriptomes.

43 proteomes composed of comparable numbers of protein domains (*i.e.* a minimum of 9,000 domains) (Fig. S8) were used to define the core-, accessory- and pan-proteomes. The core-proteome corresponds to the CCs composed of sequences from every single proteome considered, whereas the accessory-proteome corresponds to the CCs composed of sequences from a single proteome. The pan-proteome corresponds to the total number of CCs identified in the network. In addition to the Interproscan annotation process, sequences belonging to core CCs were compared to 3 databases: (i) BUSCO core eukaryotic gene set (Simão *et al.* 2015), (ii) the UniProtKB/Swiss-Prot database, and (iii) the nr database, using BLASTp and an e-value of 1e-25.

To further explore the composition and structure of the CCs, we computed the Pielou equitability index (Mulder *et al.* 2004), classically used in ecology in order to estimate the richness and/or evenness of species in a sample. Here the Pielou index was used to estimate the contribution of each proteome in any given CC, for instance for assessing whether a CC is mainly composed of domains from a limited number of proteomes. The index ranges from 0 to 1, the more homogeneous the composition of a CC, the higher the index.

### Investigation of the functional annotation

Analyses of functional traits were based on the SSN encompassing the 46 proteomes derived from “high quality” transcriptomes. The information about 10 selected functional traits was retrieved from the literature (Tab. 1). The details about plastid origin and presence were retrieved from (Caruana & Malin 2014). Dinoflagellates that are capable of mixotrophy were listed in (Jeong *et al.* 2010). The information on species harmful to humans (AZP, DSP, NSP, PSP, CFP syndromes) or to marine fauna (ichyotoxicity) was obtained from the Taxonomic Reference List of Harmful MicroAlgae of the IOC-UNESCO (http://www.marinespecies.org/hab/index.php). Dinoflagellate plastidy is reviewed in (Gagat *et al.* 2014). Dinoflagellates which have the capacity to produce DMSP in high cellular concentration were described in (Caruana *et al.* 2012). Presence of the theca, characteristic of thecate dinoflagellates, has been studied in (Lin 2011; Orr *et al.* 2012). In (Rengefors *et al.* 1998) authors studied dinoflagellates species that go through a cyst stage during their life cycle. Symbiotic taxa are characterized in (Trench & Blank 1987; Siano *et al.* 2010; Decelle *et al.* 2012; Probert *et al.* 2014; Yuasa *et al.* 2016). We later focused on CCs that are specific to a given trait, called “trait-CCs”, defined by CCs exclusively composed of vertices tagged with this trait (and excluding <NA> tags).

Following an exploratory approach, among trait-CCs, CCs including a maximum of distinct proteomes were sought (except for the “parasite” trait, as only one of the two proteomes are represented in the network). In this study, we examined more specifically the functional composition for the “harmful for human” and “symbiosis” trait-CCs. To validate the SSN capacity to detect trait-CCs characteristic for a given function, we followed a knowledge-based approach searching for sequence similarities through BLASTp (e-value 1e-3) to well-known genes from the literature.

### Research of toxin sequences in “harmful for human” trait-CCs

Specific studies on toxic dinoflagellate species have led to the establishment of defined gene sets related to toxin production that can be used as a reference for knowledge-based approaches (Snyder *et al.*; Monroe & Van Dolah 2008; Wang 2008; Sheng *et al.* 2010; Kellmann *et al.* 2010; Stüken *et al.* 2011; Salcedo *et al.* 2012; Hackett *et al.* 2013; Cusick & Sayler 2013; Lehnert *et al.* 2014; Perini *et al.* 2014; Zhang *et al.* 2014; Kohli *et al.* 2015, 2016; Meyer *et al.* 2015; Murray *et al.* 2015; Beedessee *et al.* 2015). Many of the toxic metabolites produced by some dinoflagellate species are of polyketide origin (Kellmann *et al.* 2010). 2,632 polyketide synthase (PKS) peptide sequences from (Kohli *et al.* 2016) (supplementary data 3) were compared to sequences from “harmful for human” trait-CCs to unveil PKS presence as well as non-“harmful for human” trait-CCs as a control (retained alignments show 80% sequence identity and 80% sequence coverage). Previous studies have also identified *sxt* genes involved in saxitoxin (STX) biosynthesis (Orr *et al.* 2013) that we compared to “harmful for human” and non-“harmful for human” trait-CCs using a specific threshold (retained alignments with 80% sequence identity and 90% sequence coverage). Of these, two genes have been highlighted to be related with the STX biosynthesis pathway: *sxtA* and *sxtG*. We based our investigations on 117 *sxtA1-4* and 20 *sxtG* sequences from (Murray *et al.* 2015) (Tab. S20). The differential composition of functional annotations between “harmful for human” and non-“harmful for human” trait-CCs was investigated to detect functions that are likely more represented in toxic species. The counts of each annotation found in each functional category were respectively normalized by the total number of sequences that composed both trait-CCs. Finally, the difference of pair normalized counts for the same annotation in “harmful for human” and non-“ harmful for human” trait-CCs was calculated

### Probing “symbiosis” trait-CCs

In this study, three additional transcriptomes of symbiotic species were added to the MMETSP data to increase the number of transcriptomes of symbiotic species from 10 to 13. Following a similar strategy as for the “harmful for human” CC-trait, investigation of the “symbiosis” trait in our network was based on reported sets of genes potentially involved in the symbiotic lifestyle for *Symbiodinium kawaguti* (Lin *et al.* 2015) and coral symbiotic relationships (Tab. S21). We combined this set with other putative proteins highly up-regulated in anemone-dinoflagellate symbiosis (Lehnert *et al.* 2014). The distribution of 150 “symbiotic” marker sequences was studied across “symbiosis” trait-CCs (Tab. S15). The differential composition of functional annotations between “symbiosis” and non-“symbiosis” trait-CCs was investigated as previously described for “harmful for human” trait-CCs.

## DATA ACCESSIBILITY

Link to data: http://application.sb-roscoff.fr/project/radiolaria/

## Acknowledgments

We thank Gaëlle Lelandais and Éric Pelletier for their support and critical discussions. We are also grateful to RCC staff for providing dinoflagellate cultures as well as ABIMs staff for the help on computational facilities. This work was supported by a 3-year Ph.D. grant from “Interface Pour le Vivant” (IPV) program at the Université Pierre et Marie Curie (UPMC), Paris. This project was supported by grants from Région Ile-de-France.

## REFERENCES

Altschul SF, Gish W, Miller W, Myers EW, Lipman DJ (1990) Basic local alignment search tool. Journal of Molecular Biology, 215, 403–410.

Alvarez-Ponce D, Lopez P, Bapteste E, McInerney JO (2013) Gene similarity networks provide tools for understanding eukaryote origins and evolution. Proceedings of the National Academy of Sciences of the United States of America, 110, E1594–1603.

Aranda M, Li Y, Liew YJ et al. (2016) Genomes of coral dinoflagellate symbionts highlight evolutionary adaptations conducive to a symbiotic lifestyle. Scientific Reports, 6.

Armengaud J, Trapp J, Pible O et al. (2014) Non-model organisms, a species endangered by proteogenomics. Journal of Proteomics, 105, 5–18.

Atkinson HJ, Morris JH, Ferrin TE, Babbitt PC (2009) Using Sequence Similarity Networks for Visualization of Relationships Across Diverse Protein Superfamilies. PLoS ONE, 4.

Bachvaroff TR, Gornik SG, Concepcion GT et al. (2014) Dinoflagellate phylogeny revisited: Using ribosomal proteins to resolve deep branching dinoflagellate clades. Molecular Phylogenetics and Evolution, 70, 314–322.

Beedessee G, Hisata K, Roy MC, Satoh N, Shoguchi E (2015) Multifunctional polyketide synthase genes identified by genomic survey of the symbiotic dinoflagellate, Symbiodinium minutum. BMC Genomics, 16.

Bittner L, Halary S, Payri C et al. (2010) Some considerations for analyzing biodiversity using integrative metagenomics and gene networks. Biology Direct, 5, 47.

Brocchieri L, Karlin S (2005) Protein length in eukaryotic and prokaryotic proteomes. Nucleic Acids Research, 33, 3390–3400.

Buchfink B, Xie C, Huson DH (2015) Fast and sensitive protein alignment using DIAMOND. Nature Methods, 12, 59–60.

Caron DA, Alexander H, Allen AE et al. (2016) Probing the evolution, ecology and physiology of marine protists using transcriptomics. Nature Reviews Microbiology, advance online publication.

Caruana AMN, Malin G (2014) The variability in DMSP content and DMSP lyase activity in marine dinoflagellates. Progress in Oceanography, 120, 410–424.

Caruana AMN, Steinke M, Turner SM, Malin G (2012) Concentrations of dimethylsulphoniopropionate and activities of dimethylsulphide-producing enzymes in batch cultures of nine dinoflagellate species. Biogeochemistry, 110, 87–107.

Charpentier M, Bredemeier R, Wanner G et al. (2008) Lotus japonicus CASTOR and POLLUX Are Ion Channels Essential for Perinuclear Calcium Spiking in Legume Root Endosymbiosis. The Plant Cell, 20, 3467–3479.

Cheng S, Karkar S, Bapteste E et al. (2014) Sequence similarity network reveals the imprints of major diversification events in the evolution of microbial life. Frontiers in Ecology and Evolution, 2.

Csárdi G, Nepusz T (2006) The igraph software package for complex network research. InterJournal Complex Systems.

Cusick KD, Sayler GS (2013) An Overview on the Marine Neurotoxin, Saxitoxin: Genetics, Molecular Targets, Methods of Detection and Ecological Functions. Marine Drugs, 11, 991–1018.

Day EK, Sosale NG, Lazzara MJ (2016) Cell signaling regulation by protein phosphorylation: a multivariate, heterogeneous, and context-dependent process. Current Opinion in Biotechnology, 40, 185–192.

Decelle J, Colin S, Foster RA (2015) Photosymbiosis in Marine Planktonic Protists. In: Marine Protists (eds Ohtsuka S, Suzaki T, Horiguchi T, Suzuki N, Not F), pp. 465–500. Springer Japan.

Decelle J, Probert I, Bittner L et al. (2012) An original mode of symbiosis in open ocean plankton. Proceedings of the National Academy of Sciences, 109, 18000–18005.

Dupont CL, McCrow JP, Valas R et al. (2015) Genomes and gene expression across light and productivity gradients in eastern subtropical Pacific microbial communities. The ISME Journal, 9, 1076–1092.

Flewelling LJ, Naar JP, Abbott JP et al. (2005) Brevetoxicosis: Red tides and marine mammal mortalities. Nature, 435, 755–756.

Forster D, Bittner L, Karkar S et al. (2015) Testing ecological theories with sequence similarity networks: marine ciliates exhibit similar geographic dispersal patterns as multicellular organisms. BMC Biology, 13, 16.

Gagat P, Bodył A, Mackiewicz P, Stiller JW (2014) Tertiary Plastid Endosymbioses in Dinoflagellates. In: Endosymbiosis (ed Löffelhardt W), pp. 233–290. Springer Vienna.

Gast RJ, Moran DM, Dennett MR, Caron DA (2007) Kleptoplasty in an Antarctic dinoflagellate: caught in evolutionary transition? Environmental Microbiology, 9, 39–45.

Gerlt JA, Babbitt PC, Jacobson MP, Almo SC (2012) Divergent Evolution in Enolase Superfamily: Strategies for Assigning Functions. The Journal of Biological Chemistry, 287, 29–34.

Germot A, Philippe H (1999) Critical Analysis of Eukaryotic Phylogeny: A Case Study Based on the HSP70 Family. Journal of Eukaryotic Microbiology, 46, 116–124.

Goodson MS, Whitehead LF, Douglas AE (2001) Symbiotic dinoflagellates in marine Cnidaria: diversity and function. Hydrobiologia, 461, 79–82.

Grabherr MG, Haas BJ, Yassour M et al. (2011) Full-length transcriptome assembly from RNA-Seq data without a reference genome (Trinity). Nature Biotechnology, 29, 644–652.

Haas BJ, Papanicolaou A, Yassour M et al. (2013) De novo transcript sequence reconstruction from RNAseq using the Trinity platform for reference generation and analysis. Nature Protocols, 8, 1494–1512.

Hackett JD, Wisecaver JH, Brosnahan ML et al. (2013) Evolution of Saxitoxin Synthesis in Cyanobacteria and Dinoflagellates. Molecular Biology and Evolution, 30, 70–78.

Jaeckisch N, Yang I, Wohlrab S et al. (2011) Comparative Genomic and Transcriptomic Characterization of the Toxigenic Marine Dinoflagellate Alexandrium ostenfeldii. PLOS ONE, 6, e28012.

Janouškovec J, Gavelis GS, Burki F et al. (2016) Major transitions in dinoflagellate evolution unveiled by phylotranscriptomics. Proceedings of the National Academy of Sciences, 201614842.

Jeong HJ, Yoo YD, Kim JS et al. (2010) Growth, feeding and ecological roles of the mixotrophic and heterotrophic dinoflagellates in marine planktonic food webs. Ocean Science Journal, 45, 65–91.

Jones P, Binns D, Chang H-Y et al. (2014) InterProScan 5: genome-scale protein function classification. Bioinformatics (Oxford, England), 30, 1236–1240.

Keeling PJ, Burki F, Wilcox HM et al. (2014) The Marine Microbial Eukaryote Transcriptome Sequencing Project (MMETSP): Illuminating the Functional Diversity of Eukaryotic Life in the Oceans through Transcriptome Sequencing. PLOS Biol, 12, e1001889.

Keller MB, Lavori PW, Friedman B et al. (1987) The Longitudinal Interval Follow-up Evaluation. A comprehensive method for assessing outcome in prospective longitudinal studies. Archives of General Psychiatry, 44, 540–548.

Kellmann R, Stüken A, Orr RJS, Svendsen HM, Jakobsen KS (2010) Biosynthesis and Molecular Genetics of Polyketides in Marine Dinoflagellates. Marine Drugs, 8, 1011–1048.

Khosla C, Herschlag D, Cane DE, Walsh CT (2014) Assembly Line Polyketide Synthases: Mechanistic Insights and Unsolved Problems. Biochemistry, 53, 2875–2883.

Kohli GS, John U, Figueroa RI et al. (2015) Polyketide synthesis genes associated with toxin production in two species of Gambierdiscus (Dinophyceae). BMC Genomics, 16, 410.

Kohli GS, John U, Van Dolah FM, Murray SA (2016) Evolutionary distinctiveness of fatty acid and polyketide synthesis in eukaryotes. The ISME Journal.

Langmead B, Trapnell C, Pop M, Salzberg SL (2009) Ultrafast and memory-efficient alignment of short DNA sequences to the human genome. Genome Biology, 10, R25.

Le Bescot N, Mahé F, Audic S et al. (2016) Global patterns of pelagic dinoflagellate diversity across protist size classes unveiled by metabarcoding. Environmental Microbiology, 18, 609–626.

Lee R, Lai H, Malik SB et al. (2014) Analysis of EST data of the marine protist Oxyrrhis marina, an emerging model for alveolate biology and evolution. BMC Genomics, 15, 122.

Lehnert EM, Mouchka ME, Burriesci MS et al. (2014) Extensive Differences in Gene Expression Between Symbiotic and Aposymbiotic Cnidarians. G3: Genes|Genomes|Genetics, 4, 277–295.

Lima-Mendez G, Faust K, Henry N et al. (2015) Determinants of community structure in the global plankton interactome. Science, 348, 1262073.

Lin S (2011) Genomic understanding of dinoflagellates. Research in Microbiology, 162, 551–569.

Lin S, Cheng S, Song B et al. (2015) The Symbiodinium kawagutii genome illuminates dinoflagellate gene expression and coral symbiosis. Science, 350, 691–694.

Lionetti V, Metraux J-P (2015) Plant cell wall in pathogenesis, parasitism and symbiosis. Frontiers Media SA.

Lopez P, Halary S, Bapteste E (2015) Highly divergent ancient gene families in metagenomic samples are compatible with additional divisions of life. Biology Direct, 10, 64.

Marchler-Bauer A, Anderson JB, Cherukuri PF et al. (2005) CDD: a Conserved Domain Database for protein classification. Nucleic Acids Research, 33, D192–D196.

Massana R, Gobet A, Audic S et al. (2015) Marine protist diversity in European coastal waters and sediments as revealed by high-throughput sequencing. Environmental Microbiology, 17, 4035–4049.

Matzke M, Weiger TM, Papp I, Matzke AJM (2009) Nuclear membrane ion channels mediate root nodule development. Trends in Plant Science, 14, 295–298.

Méheust R, Zelzion E, Bhattacharya D, Lopez P, Bapteste E (2016) Protein networks identify novel symbiogenetic genes resulting from plastid endosymbiosis. Proceedings of the National Academy of Sciences, 113, 3579–3584.

Meyer JM, RöWdelsperger C, Eichholz K et al. (2015) Transcriptomic characterisation and genomic glimps into the toxigenic dinoflagellate Azadinium spinosum, with emphasis on polykeitde synthase genes. BMC Genomics, 16.

Monroe EA, Van Dolah FM (2008) The Toxic Dinoflagellate Karenia brevis Encodes Novel Type I-like Polyketide Synthases Containing Discrete Catalytic Domains. Protist, 159, 471–482.

Montagnes DJS, Lowe CD, Roberts EC et al. (2011) An introduction to the special issue: Oxyrrhis marina, a model organism? Journal of Plankton Research, 33, 549–554.

Mulder CPH, Bazeley-White E, Dimitrakopoulos PG et al. (2004) Species evenness and productivity in experimental plant communities. Oikos, 107, 50–63.

Murray SA, Diwan R, Orr RJS, Kohli GS, John U (2015) Gene duplication, loss and selection in the evolution of saxitoxin biosynthesis in alveolates. Molecular Phylogenetics and Evolution, 92, 165–180.

Murray SA, Suggett DJ, Doblin MA et al. (2016) Unravelling the functional genetics of dinoflagellates: a review of approaches and opportunities. Perspectives in Phycology, 37–52.

Orr RJS, Murray SA, Stüken A, Rhodes L, Jakobsen KS (2012) When Naked Became Armored: An Eight-Gene Phylogeny Reveals Monophyletic Origin of Theca in Dinoflagellates (S Lin, Ed,). PLoS ONE, 7, e50004.

Orr RJS, Stüken A, Murray SA, Jakobsen KS (2013) Evolutionary Acquisition and Loss of Saxitoxin Biosynthesis in Dinoflagellates: the Second `Core Gene, sxtG. Applied and Environmental Microbiology, 79, 2128–2136.

Parra G, Bradnam K, Korf I (2007) CEGMA: a pipeline to accurately annotate core genes in eukaryotic genomes. Bioinformatics (Oxford, England), 23, 1061–1067.

Perini F, Galluzzi L, Dell'Aversano C et al. (2014) SxtA and sxtG Gene Expression and Toxin Production in the Mediterranean Alexandrium minutum (Dinophyceae). Marine Drugs, 12, 5258–5276.

Probert I, Siano R, Poirier C et al. (2014) Brandtodinium gen. nov. and B. nutricula comb. Nov. (Dinophyceae), a dinoflagellate commonly found in symbiosis with polycystine radiolarians. Journal of Phycology, 50, 388–399.

Rengefors K, Karlsson I, Hansson L-A (1998) Algal cyst dormancy: a temporal escape from herbivory. Proceedings of the Royal Society B: Biological Sciences, 265, 1353–1358.

Salcedo T, Upadhyay RJ, Nagasaki K, Bhattacharya D (2012) Dozens of Toxin-Related Genes Are Expressed in a Nontoxic Strain of the Dinoflagellate Heterocapsa circularisquama. Molecular Biology and Evolution, 29, 1503–1506.

Sheng J, Malkiel E, Katz J, Adolf JE, Place AR (2010) A dinoflagellate exploits toxins to immobilize prey prior to ingestion. Proceedings of the National Academy of Sciences, 107, 2082–2087.

Shoguchi E, Shinzato C, Kawashima T et al. (2013) Draft Assembly of the Symbiodinium minutum Nuclear Genome Reveals Dinoflagellate Gene Structure. Current Biology, 23, 1399–1408.

Siano R, Alves-de-Souza C, Foulon E et al. (2011) Distribution and host diversity of Amoebophryidae parasites across oligotrophic waters of the Mediterranean Sea. Biogeosciences, 8, 267–278.

Siano R, Montresor M, Probert I, Not F, de Vargas C (2010) Pelagodinium gen. nov. and P. bUii comb. nov., a dinoflagellate symbiont of planktonic foraminifera. Protist, 161, 385–399.

Sibbald SJ, Archibald JM (2017) More protist genomes needed. Nature Ecology & Evolution, 1, 145.

Simão FA, Waterhouse RM, Ioannidis P, Kriventseva EV, Zdobnov EM (2015) BUSCO: assessing genome assembly and annotation completeness with single-copy orthologs. Bioinformatics, btv351.

Snyder RV, Gibbs PDL, Palacios A et al. Polyketide Synthase Genes from Marine Dinoflagellates. Marine Biotechnology, 5, 1–12.

Stoecker DK, Hansen PJ, Caron DA, Mitra A (2017) Mixotrophy in the Marine Plankton. Annual Review of Marine Science, 9, 311–335.

Stüken A, Orr RJS, Kellmann R et al. (2011) Discovery of Nuclear-Encoded Genes for the Neurotoxin Saxitoxin in Dinoflagellates. PLOS ONE, 6, e20096.

Trench RK, Blank RJ (1987) Symbiodinium Microadriaticum Freudenthal, S. Goreauii Sp. Nov., S. Kawagutii Sp. Nov. and S. Pilosum Sp. Nov.: Gymnodinioid Dinoflagellate Symbionts of Marine Invertebrates 1. Journal of Phycology, 23, 469–481.

Vonk FJ, Casewell NR, Henkel CV et al. (2013) The king cobra genome reveals dynamic gene evolution and adaptation in the snake venom system. Proceedings of the National Academy of Sciences, 110, 20651–20656.

Wang D-Z (2008) Neurotoxins from Marine Dinoflagellates: A Brief Review. Marine Drugs, 6, 349–371.

Yang Y, Smith SA (2013) Optimizing de novo assembly of short-read RNA-seq data for phylogenomics. BMC Genomics, 14, 328.

Yang L, Tan J, O'Brien EJ et al. (2015) Systems biology definition of the core proteome of metabolism and expression is consistent with high-throughput data. Proceedings of the National Academy of Sciences, 112, 10810–10815.

Yuasa T, Horiguchi T, Mayama S, Takahashi O (2016) Gymnoxanthella radiolariae gen. et sp. nov. (Dinophyceae), a dinoflagellate symbiont from solitary polycystine radiolarians. Journal of Phycology, 52, 89–104.

Zhang Y, Zhang S-F, Lin L, Wang D-Z (2014) Comparative Transcriptome Analysis of a Toxin-Producing Dinoflagellate Alexandrium catenella and Its Non-Toxic Mutant. Marine Drugs, 12, 5698–5718.

